# *cfrB, cfrC*, and a potential new *cfr*-like gene in *Clostridium difficile* strains recovered across Latin America

**DOI:** 10.1101/649020

**Authors:** Vanja Stojković, María Fernanda Ulate, Fanny Hidalgo-Villeda, Emmanuel Aguilar, Camilo Monge-Cascante, Marjorie Pizarro-Guajardo, Kaitlyn Tsai, Edgardo Tzoc, Margarita Camorlinga, Daniel Paredes-Sabja, Carlos Quesada-Gómez, Danica Galonić Fujimori, César Rodríguez

## Abstract

Cfr is a radical *S*-adenosyl-L-methionine (SAM) enzyme that confers cross-resistance to all antibiotics targeting the large ribosomal subunit through hypermethylation of nucleotide A2503 of 23S rRNA. Of the four known *cfr* genes known to date, *cfr*(B) and *cfr*(C) have been sporadically found in *C. difficile*, yet functional characterization of *cfr*(C) is still lacking. We identified genes for putative Cfr-like enzymes among clinical *C. difficile* strains from Mexico, Honduras, Costa Rica, and Chile. To confirm their identity and activity, we obtained minimum inhibitory concentrations for ribosome-targeting antibiotics, annotated whole genome sequences, and performed a functional characterization of Cfr(C). The seven representative isolates analyzed displayed different levels of resistance to PhLOPS_A_ antibiotics in the absence of the ribosome protection factor OptrA, and mutations in genes for 23S rRNAs or the ribosomal proteins L3 and L4. *cfr*(B) was detected in four isolates as part of a Tn*6218*-like transposon or an un-described mobile genetic element. In turn, *cfr*(C) was found integrated into an ICE-element. One isolate harbored a putative *cfr*-like gene that shows only 51-58% of sequence identity to Cfr and known Cfr-like enzymes. Moreover, our *in vitro* assays confirmed that Cfr(C) methylates *E. coli and C. difficile* 23S rRNA fragments. These results indicate selection of *cfr*-like genes in *C. difficile* from Latin America, suggest that the diversity of *cfr*-like resistance genes is larger than anticipated, and provide the first assessment of the methylation activity of Cfr(C).

## 1. Introduction

The bacterial ribosome is one of the most common targets for antibiotics of clinical and veterinary relevance. Resistance to ribosome-targeting antibiotics occurs primarily through modification of drug’s binding sites, specifically through mutation or modification of ribosomal RNA (rRNA) or ribosomal proteins ^1^. Several rRNA modifying enzymes implicated in antibiotic resistance have been discovered ^2^, and among them, the radical SAM enzyme Cfr is noteworthy because it provides cross-resistance to <u>P</u>henicols (e.g. thiamphenicol), <u>L</u>incosamides (e.g. clindamycin), <u>O</u>xazolidinones (e.g. linezolid), <u>P</u>leuromutilins (e.g. tiamulin), and <u>S</u>treptogramin A (e.g. dalfopristin) through C8 methylation of the A2503 residue in 23S rRNA (*E. coli* numbering), which is located in the peptidyl transferase center (PTC) ^3^. In addition to this so-called PhLOPS_A_ phenotype ^4^, Cfr-mediated methylation leads to resistance to 16-member macrolides, the aminocyclitol hygromycin A, and the nucleoside antimicrobial agent A201A ^4–6^.

*cfr* and *cfr*-like genes are typically found on mobile genetic elements (MGEs). Moreover, since acquisition of *cfr* exhibits low fitness costs^7^, the spread of these resistance genes threatens the utility of PTC-targeting antibiotics in the clinic. The *cfr* gene was first discovered on a *Staphylococcus sciuri* plasmid ^8^, but it is now found in nearly twenty different molecular contexts in isolates of *Enterococcus, Bacillus, Proteus vulgaris, Escherichia coli, Macrococcus caseolyticus, Jeotgalicoccus pinnipedialis*, and *Streptoccocus suis* from Europe, Latin America, USA, and Asia ^3^. Homologues of *cfr* have been identified in non-pathogenic Bacillales ^9^ and three additional *cfr*-like genes sharing less than 80% protein sequence identity to Cfr have been described in *Clostridium* and *Enterococcus* [3]. These genes are known as *cfr*(B), *cfr*(C), and c*fr*(D).

In *C. difficile, cfr*(B) was first detected in strain 11140508 contained within Tn*6218*-like elements ^10,11^. Afterwards, Candela *et al.* defined *cfr*(C) after analysis of *C. difficile* T10 and found it in three types of integrative and conjugative elements (ICEs) in several other strains, including the non-toxigenic strain *C. difficile* F548^12^. Subsequently, Hansen and Vester demonstrated by primer extension that a codon-optimized version of *cfr*(B) of *C. difficile* 11140508 modifies A2503 in 23S rRNA when expressed in *E. coli* ^13^. Equivalent functional evidence is missing for Cfr(C), though it has been shown to confer PhLOPS_A_ resistance upon introduction into the linezolid-susceptible strain *C. difficile* 630Δ*erm* ^12^.

Linezolid is not used to treat *C. difficile* infections (CDI) despite its confirmed utility to prevent CDI in patients with ventilator associated pneumonia ^14^ and to reduce *C. difficile* toxin gut levels in a mice model ^15^. Moreover, the closely related antibiotic cadazolid inhibits moxifloxacin-resistant *C. difficile* NAP1/027 strains without affecting gut commensals, ^16^ and though it did not pass a Phase III trial ^17^, novel oxazolidinones to treat *C. difficile* infections may appear in the future.

Based on this potential utility of oxazolidinones in treating CDI and the current use of linezolid to treat infections caused by anaerobic bacteria in Mexico and Honduras (personal communication), we investigated seven clinical *C. difficile* isolates from Latin America that circulated between 2009 and 2016 to determine whether they carry functional *cfr* or *cfr*-like genes. To this end, we obtained minimum inhibitory concentrations (MICs) for several PTC-targeting antibiotics, analyzed draft whole genome sequences (WGS), and evaluated the *in vitro* activity of the Cfr(C) enzyme detected in two clinical isolates.

## 2. Methods

### 2.1. Strains

This study included ribotype- or PFGE-confirmed NAP1/027/ST01 clinical isolates from Mexico (DF11), Honduras (HON06, HON10, HON11) and Chile (PUC51, PUC347), and one isolate from the NAP_CR1_/012/ST54 genotype from Costa Rica (LIBA5707). These bacteria were recovered between 2009-2016 from stool samples of human patients and represent strains that were shown by automated annotation of WGS to carry sequences for potential Cfr enzymes (^18,19^, and unpublished data). With a single exception (DF11, recovered from a 3-years old patient), all isolates were obtained from adults with active diarrhea compatible with CDI. DF, PUC, and LIBA isolates were obtained during confirmed CDI outbreaks. The linezolid-susceptible strain LIBA5701 was used as a negative control in the determinations of minimum inhibitory concentrations (MIC) because it is a NAP_CR1_ strain that lacks *cfr*-like genes (see below) ^18^.

### 2.2. MIC determinations

MIC of clindamycin and linezolid were obtained for isolates from México (DF), Honduras (HON) and Costa Rica (LIBA) using E-test strips containing a 0.16 to 256 µg/ml concentration gradient (BioMerieux). This isolate subset was also analyzed by agar microdilution ^20^ using brain heart infusion plates containing 1-256 µg/ml of tiamulin or thiamphenicol. The susceptibility of the Chilean isolates (PUC) to linezolid, tiamulin, and thiamphenicol was assessed using agar macrodilution with brain heart infusion plates containing 1-256 µg/ml of the corresponding antibiotics. *C. difficile* ATCC 70057 (linezolid^s^) was tested in parallel for quality control purposes.

### 2.3. Comparative genomics

WGS were obtained by sequencing-by-synthesis using multiplexed paired-end libraries and Illumina HiSeq2000 or Miseq platforms. After trimming with sickle (https://github.com/najoshi/sickle), reads were assembled using Spades v.3.12 ^21^. For automated annotation we used Prokka v. 1.13 ^22^. Regions of interest were trimmed and reannotated using BLAST, BLASTP, eggNOG ^23^ and UniProt searches. Resistance genes were identified manually or with the CARD database v.3.0.1 ^24^. Megablast searches against the NCBI *nr* database were used to identify sequences resembling the MGEs here defined. All genomes and genome comparisons were visualized in Artemis or ACT, respectively. Linear comparison figures were prepared with Easyfig. The presence of SNPs or indels in genes encoding linezolid binding sites ^25^, including genes from 23 rRNAs and the ribosomal proteins L3 and L4, was checked in all isolates through bwa mapping of trimmed reads to WGS from the reference strains R20291 (accession number FN545816) or CD630 (accession number AM180355). Trimmed reads and assemblies for the DF and HON isolates can be downloaded from the MicrobesNG portal (https://microbesng.uk/portal/projects/405FF6AC-A5E0-E04A-AECF-A5C9371B8B60/). Sequencing data for LIBA5707 is available at the European Nucleotide Archive (run ERR467555). Data for PUC51 and PUC347 can be retrieved using the accession numbers ERZ816937 and ERZ816944, respectively.

### 2.4. Comparison of RlmN and Cfr protein sequences

Though both RlmN and Cfr modify A2503, the former is a housekeeping gene and the latter an acquired antibiotic resistance gene ^26^. To examine the relationship between putative Cfr sequences mentioned in this paper to other Cfr and RlmN sequences, we performed a phylogenetic analysis. To this end, Cfr-like and RlmN-like orthologs from selected Firmicutes species (Supplementary Table 2) were retrieved from the Integrated Microbial Genomes-Joint Genome institute (IMG/JGI) database by BLAST using the RlmN sequence from *Bacillus subtilis* as a query, as done in ^27^. Additional RlmN/Cfr paralogous sequences from *Paenibacillus durus* were retrieved from the NCBI. These sequences were aligned using MUSCLE ^28^, prior to the generation of phylogenetic tree by PhyML with the Akaike Information Criterion for model selection ^29^.

**Table 2.**
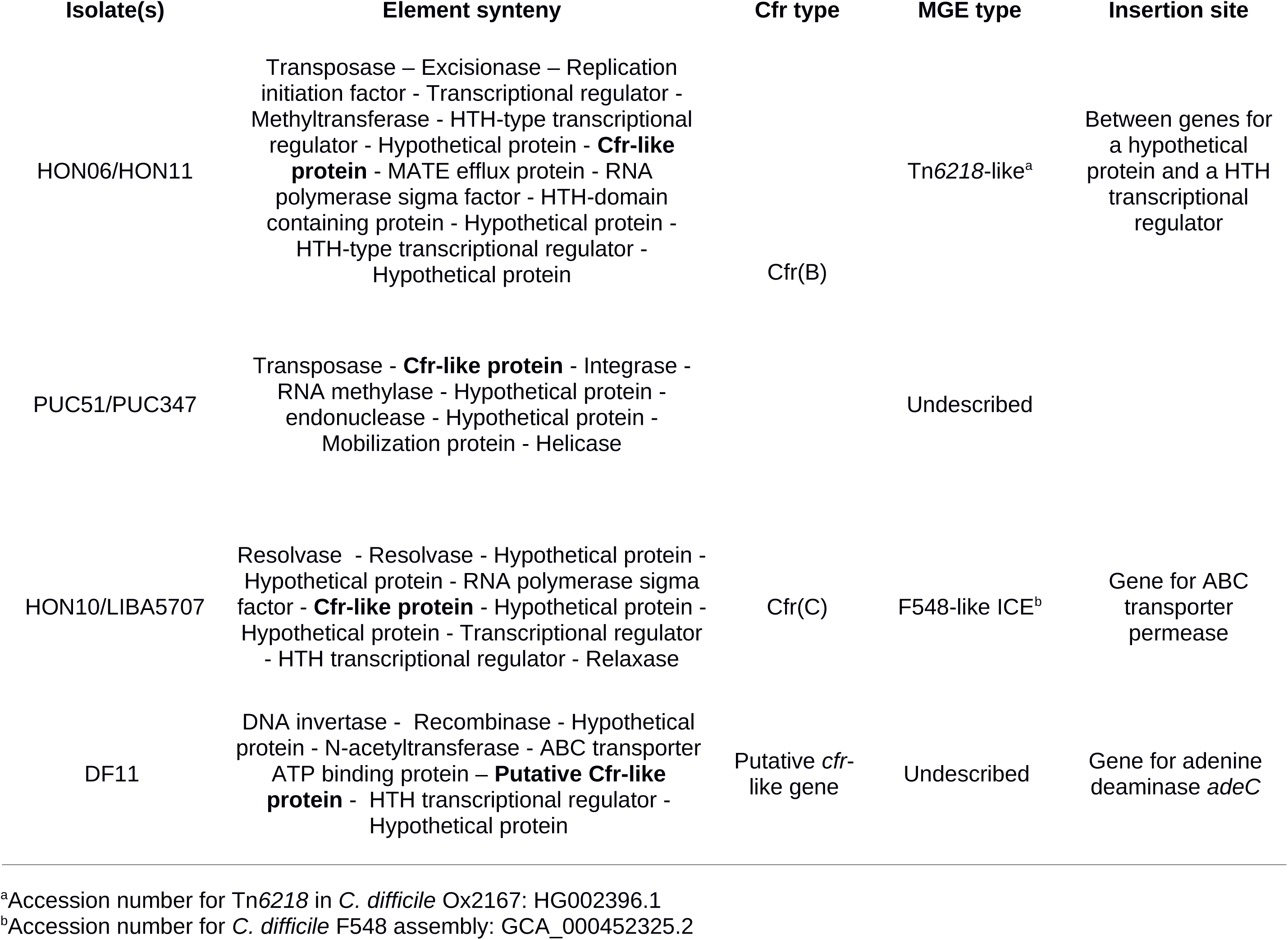
Annotation of the putative mobile genetic elements (MGE) in which *cfr*-like genes were detected

| Isolate(s) | Element synteny | Cfr type | MGE type | Insertion site |
| --- | --- | --- | --- | --- |
| HON06/HON11 | Transposase – Excisionase – Replication initiation factor - Transcriptional regulator - Methyltransferase - HTH-type transcriptional regulator - Hypothetical protein - **Cfr-like protein** - MATE efflux protein - RNA polymerase sigma factor - HTH-domain containing protein - Hypothetical protein - HTH-type transcriptional regulator - Hypothetical protein | Cfr(B) | Tn6218-like<sup>a</sup> | Between genes for a hypothetical protein and a HTH transcriptional regulator |
| PUC51/PUC347 | Transposase - **Cfr-like protein** - Integrase - RNA methylase - Hypothetical protein - endonuclease - Hypothetical protein - Mobilization protein - Helicase |  | Undescribed |  |
| HON10/LIBA5707 | Resolvase - Resolvase - Hypothetical protein - Hypothetical protein - RNA polymerase sigma factor - **Cfr-like protein** - Hypothetical protein - Hypothetical protein - Transcriptional regulator - HTH transcriptional regulator - Relaxase | Cfr(C) | F548-like ICE<sup>b</sup> | Gene for ABC transporter permease |
| DF11 | DNA invertase - Recombinase - Hypothetical protein - N-acetyltransferase - ABC transporter ATP binding protein – **Putative Cfr-like protein** - HTH transcriptional regulator - Hypothetical protein | Putative *cfr*-like gene | Undescribed | Gene for adenine deaminase *adeC* |

<sup>a</sup>Accession number for Tn6218 in *C. difficile* Ox2167: HG002396.1
<sup>b</sup>Accession number for *C. difficile* F548 assembly: GCA\_000452325.2

### 2.5. Expression and purification of Cfr(C)

A codon optimized sequence of *cfr(C)* was cloned into the pET21a vector and overexpressed in *Escherichia coli* BL21-CodonPlus (DE3)-RIPL as described previously ^26,30,31^. The resulting enzyme was then purified by Talon chromatography (Clontech) and underwent iron-sulfur cluster reconstitution using previously published protocols ^26,30,31^.

### 2.6. Preparation of truncated rRNA substrates for the *in vitro* methylation assay

The *E. coli* 23S rRNA fragment 2447-2625 used in the *in vitro* methylation assay (see below) was generated by *in vitro* transcription following a previously published protocol ^26,30^. *C. difficile* 23S rRNA fragments 2451-2629 and 2022-2629 were generated by *in vitro* transcription in the same manner but using different PCR products as templates. Briefly, forward PCR primers contained the T7 RNA polymerase promoter sequence TAATACGACTCACTATAGG, followed by several nucleotides corresponding to specific region of *C. difficile* 23S rRNA. 23S rRNA fragments were amplified using genomic *C. difficile* DNA purchased from the American Type Culture Collection as template.

### 2.7. *In vitro* methylation assay

*In vitro* reactions were performed in 100 µL volumes under the following conditions: 100 mM HEPES pH 8.0, 100 mM KCl, 10 mM MgCl_2_, 2 mM DTT, 20 μM Flavodoxin, 2 μM Flavodoxin reductase, 4 μM RNA and 0.14 μCi [14C-methyl]-SAM (58 mCi/mmoL) and 5-10 μM purified enzyme. Reactions were initiated by addition of 1 mM NADPH (final concentration) and were allowed to proceed at 37°C for 1.5 h. The RNA was recovered from the reaction mixtures using the RNA Clean & Concentrator kit (Zymo Research) and added to vials containing Ultima Gold scintillation fluid (Perkin Elmer). The amount of radioactivity incorporated in the product was measured using a Beckman–Coulter LS6500 multipurpose scintillation counter (Fullerton, CA, USA). Each value represents the average of at least duplicate measurements, with one standard deviation (SD) indicated.

### 2.8. HPLC separation and identification of methylated adenosines

Purified, methylated rRNA from *in vitro* reactions was enzymatically digested to mononucleosides using nuclease P_1_ (Sigma-Aldrich), snake venom phosphodiesterase (Sigma-Aldrich), and alkaline phosphatase from calf intestine (New England Biolabs) as described before ^26,30^. The digested samples were separated by HPLC using a Luna analytical C18 column (10 μm, 4.6 mm × 250 mm) (Phenomenex, Torrance, CA, USA) and a previously published protocol ^26,30^. Mononucleosides and synthetic methyladenosine standards were detected by their UV absorption at 256 nm, while the ^14^C-labeled methyladenosines were detected with a Packard radiomatic 515TR flow scintillation analyzer (Perkin Elmer).

## 3. Results

### 3.1. Detection of *cfr*-like genes

Isolates HON06, HON11, PUC51, and PUC347 carry a *cfr*(B) gene identical to that of *C. difficile* 11140508 (Table 1). On the other hand, isolates HON10 and LIBA5707 have the *cfr*(C) allele previously seen in *C. difficile* T10 (Table 1). Interestingly, the genome of isolate DF11 includes a gene for a radical SAM RNA methylating enzyme that only shares 51-58% identity with *cfr, cfr*(B), *cfr*(C), and *cfr*(D) and therefore might represent a new *cfr*-like gene according to the MLS nomenclature system maintained by Dr. Marilyn Roberts (Table 1). In congruence with this classification, the predicted protein sequence of the putative *cfr*-like gene of DF11 shows homology to C8 RNA methylating enzymes deposited in the BLASTp, EggNOG, UniProt, and SFLD databases (Supplementary Table 1). Cfr(B) and Cfr(D) form a cluster with functional Cfr enzymes. By contrast, Cfr(C) and the product of the putative *cfr*-like gene from isolate DF11 appear in a group of Cfr-like proteins that clades with Cfr sequences awaiting functional characterization (Figure 1).

**Table 1.**
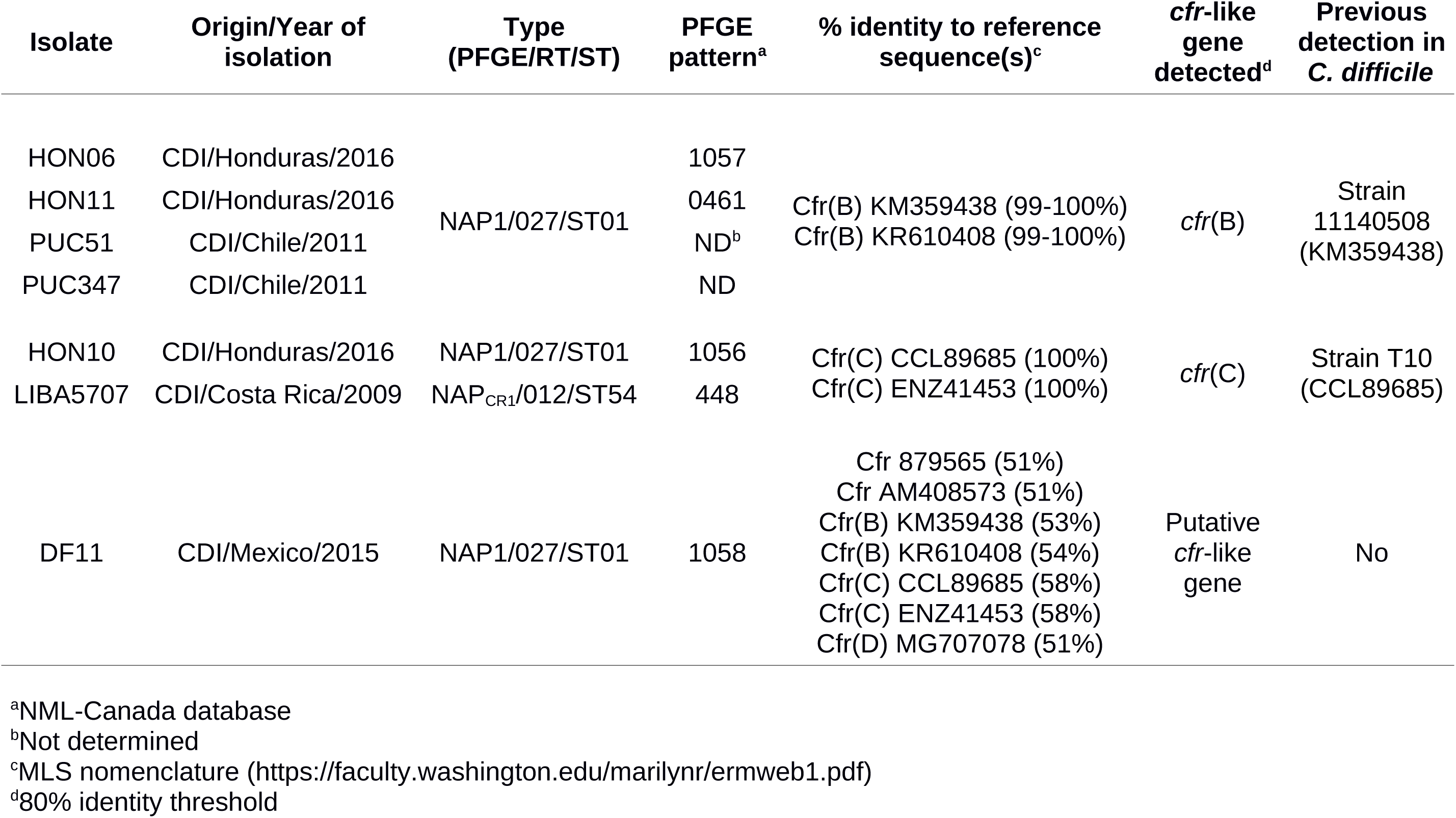
*cfr*-like genes detected

**Figure 1.**
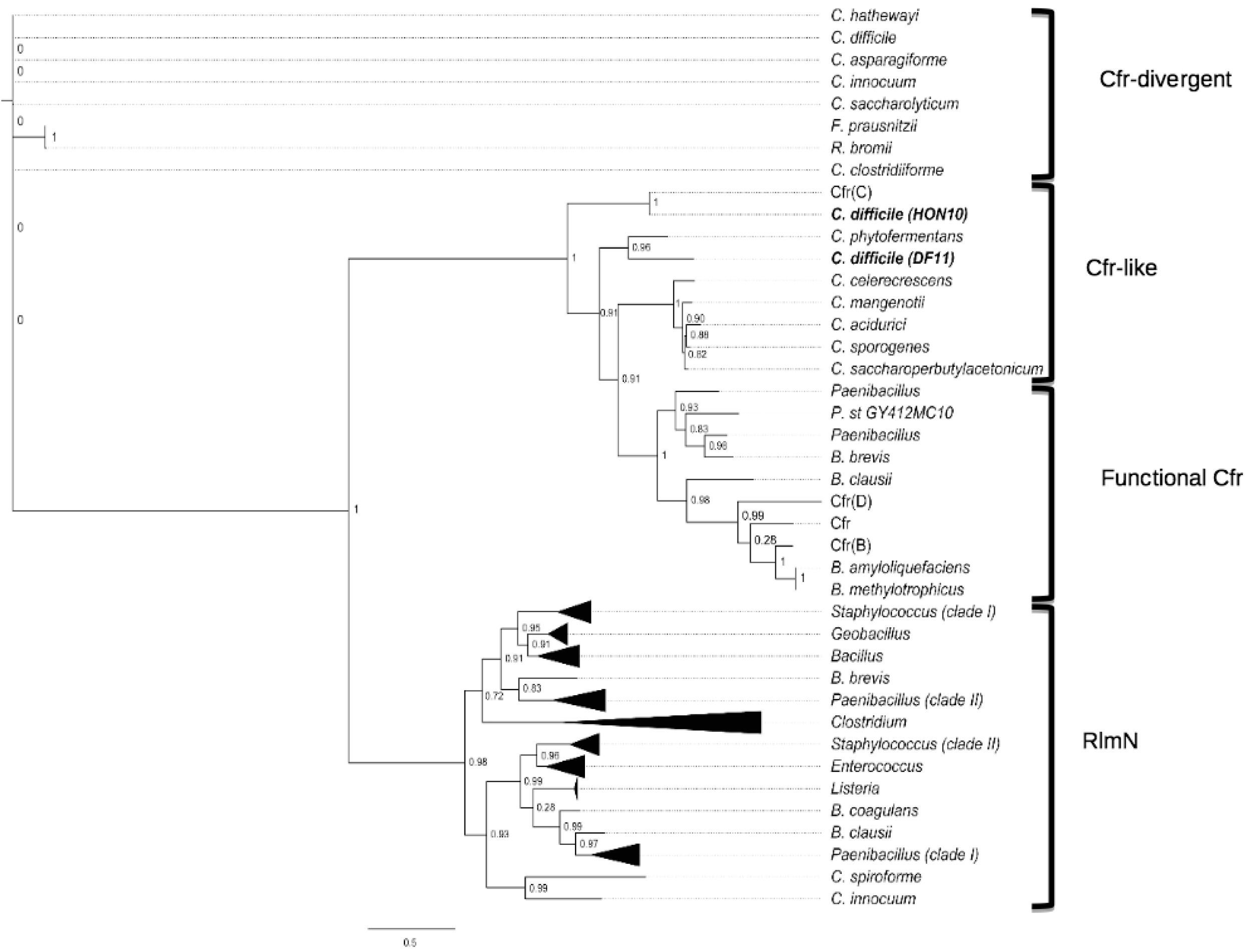
Evolutionary relationship of RlmN and Cfr sequences from selected Firmicutes species. Functionally characterized Cfrs, Cfr-like proteins, Cfr divergent proteins, and known and putative RlmNs sequences are marked. While Cfr-like proteins clade with known Cfrs lacking functional characterization, Cfr-divergent proteins diverged early in evolutionary time and do not clade with either Cfrs or RlmNs. The enzymes of isolates *C. difficile* HON10/LIBA5707 and DF11 appear highlighted in bold. The distance scale underneath the tree indicates the average number of substitutions per site. IMG/JGI database identifiers or accession numbers of protein sequences used in the tree are provided in Supplementary Table 2.

All *cfr*-like genes detected were found on four types of putative MGEs with anticipated mobilization or conjugation potential (Table 2). In detail, while isolates HON06 and HON11 have *cfr*(B) within a Tn*6218*-like element, isolates PUC51 and PUC347 have *cfr*(B) elsewhere in their genomes in an undescribed genetic structure (Table 2). The best hit for this novel MGE in a megablast search against the nr database was a genomic fragment of *Faecalibacterium prausnitzii* L2/6 (Query cover=74%, E-value=0, Identity=99%); a species that has not been previously reported to carry *cfr*-like genes. The *cfr*(C) genes of isolates HON10 and LIBA5707, in turn, were traced back to a MGE sharing similarity with the ICEs of *C. difficile* F548 ^12^ (Table 2). The putative new *cfr*-like gene of isolate DF11 was found integrated into a distinct MGE that shows partial hits to genomic sequences of various intestinal Firmicutes (Table 2), including *Lachnoclostridium* sp. YL32 (Query cover=60%, E-value=0, Identity=94%), *Roseburia intestinalis* XB6B4 (Query cover=60%, Evalue=0, Identity=92%), *Faecalibacterium prausnitzii* A2165 (Query cover=60%, Evalue=0, Identity=88%), and *C. difficile* Z31 (Query cover=60%, Evalue=0, Identity=87%). In this case, the shared regions were concentrated at the 5’and 3’ ends of the element and did not include the putative *cfr*-like gene or their immediate neighboring genes (Table 2). None of the WGS studied showed mutations or indels in 23S RNA genes, the ribosomal proteins L3 and L4, or presence of OptrA, which are mechanisms known to lead to a PhLOPS_A_ phenotype.

### 3.2. MIC

To evaluate whether the presence of *cfr*-like genes in clinical isolates leads to PhLOPS_A_ phenotype we performed susceptibility tests to several PTC-targeting antibiotics. As a negative control we used NAP_CR1_ strain LIBA5701, which lacks *cfr*-like genes and thus does not exhibit PhLOPS_A_ phenotype (Table 3). Isolates HON06 and HON11, which carry *cfr*(B) in a Tn*6218*-like context, and HON10 and LIBA5707, which is positive for *cfr*(C), exhibited an 8-24 fold increase in the MIC of linezolid and ≥128 fold increase in the MIC of tiamulin with respect to the control strain (Table 3). In a comparable manner, a 4-32 fold MIC increase with respect to the control was recorded for the same isolates when exposed to thiamphenicol (Table 3). All isolates had a MIC for clindamycin ≥256 µg/mL due to presence of the methylase ErmB (^18,19^, unpublished data). Despite carrying a *cfr*(B) gene, the MICs of linezolid, tiamulin, and thiamphenicol obtained for the Chilean isolates PUC51 and PUC347 were at least 2-4 fold lower than those obtained for the other test isolates, yet MIC of linezolid and tiamulin were still at least 12 fold higher than the MIC obtained for the LIBA5701 control (Table 3).

**Table 3.**
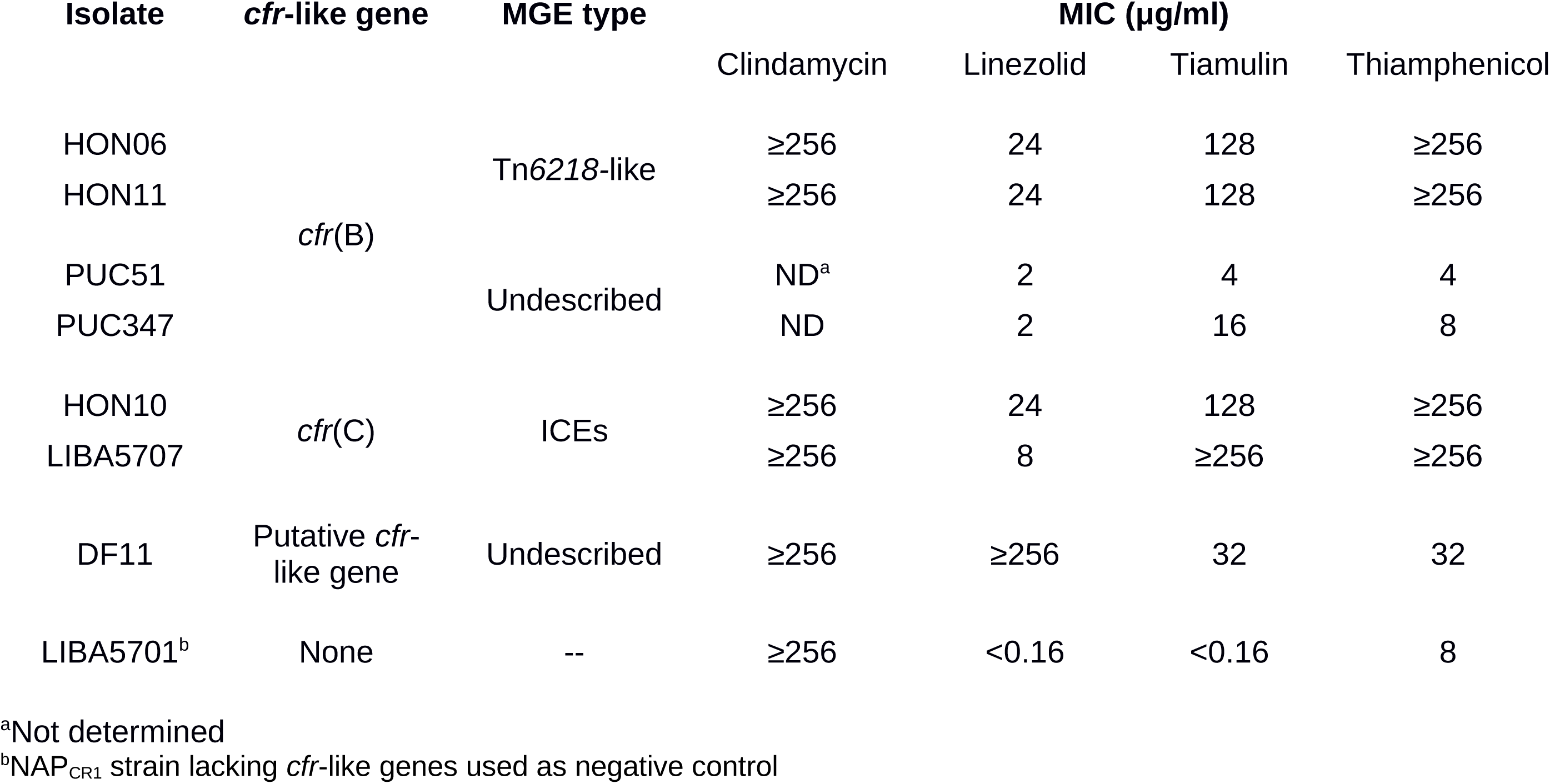
Minimum inhibitory concentrations of various PTC-targeting antibiotics

### 3.3. Functional analysis of Cfr(C)

To investigate whether Cfr(C) is a C8-methylating enzyme, we overexpressed in *Escherichia coli* a codon-optimized version of the *cfr*(C) sequence of HON10/LIBA5707. The resulting protein was purified under anaerobic conditions and its iron-sulfur cluster reconstituted. Thereafter, we performed an *in vitro* methylation assay with either *in vitro* transcribed 23S rRNA *E. coli* or *C. difficile* fragments and [^14^C-methyl]-*S*-adenosyl methionine ([^14^C-methyl]-SAM), and the amount of radioactivity incorporated into the RNA product was determined. This assay revealed that Cfr(C) can methylate *E. coli* and *C. difficile* 23S rRNA *in vitro* (Figure 2). However, while significantly above the background, the methylation level detected in the 2447-2625 rRNA fragment of *E. coli* or the 2451-2629 fragment of *C. difficile* was lower than that observed in the reaction of 23S rRNA fragments with *E. coli* RlmN (Figure 2).

**Figure 2.**
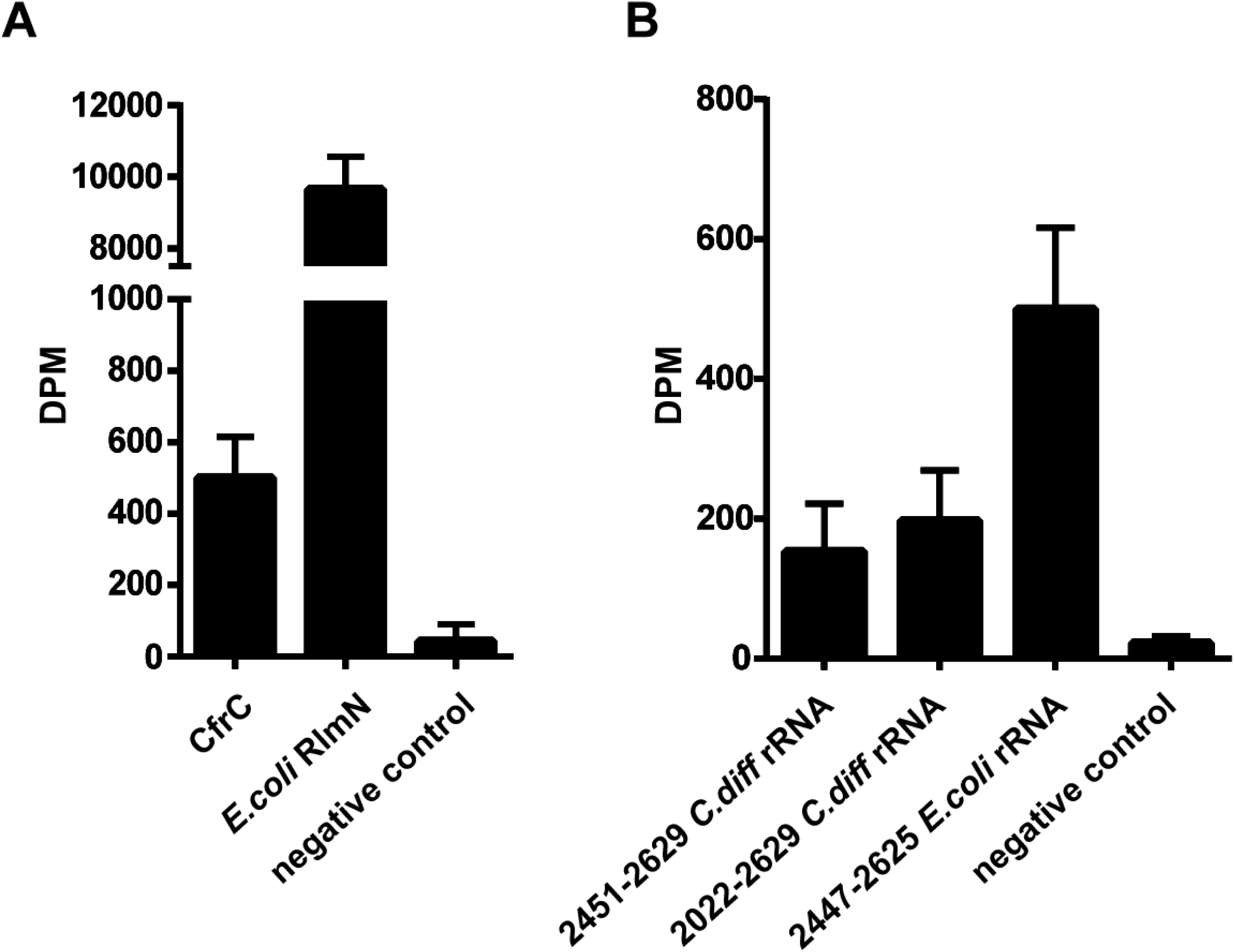
Cfr(C) and *E. coli* RlmN-mediated methylation of *in vitro* transcribed *E. coli* 2447-2625 23S rRNA fragment (A). Cfr(C)-mediated methylation of *in vitro* transcribed *E. coli* and *C. difficile* 23S rRNA fragments (B). Bars represent the mean of at least two replicates ± s.d.

To establish the regioselectivity of the modification on the adenosine ring by Cfr(C), radiolabeled RNA product isolated from the *in vitro* assay with *E. coli* RNA was purified, digested to individual nucleosides, and analyzed by HPLC. Unlike the 2-methyladenosine product of the reaction with *E. coli* RlmN, the product of the reaction with purified Cfr(C) co-eluted with the 8-methyladenosine standard, indicating that this enzyme methylates A2503 at the C8 position (Figure 3).

**Figure 3.**
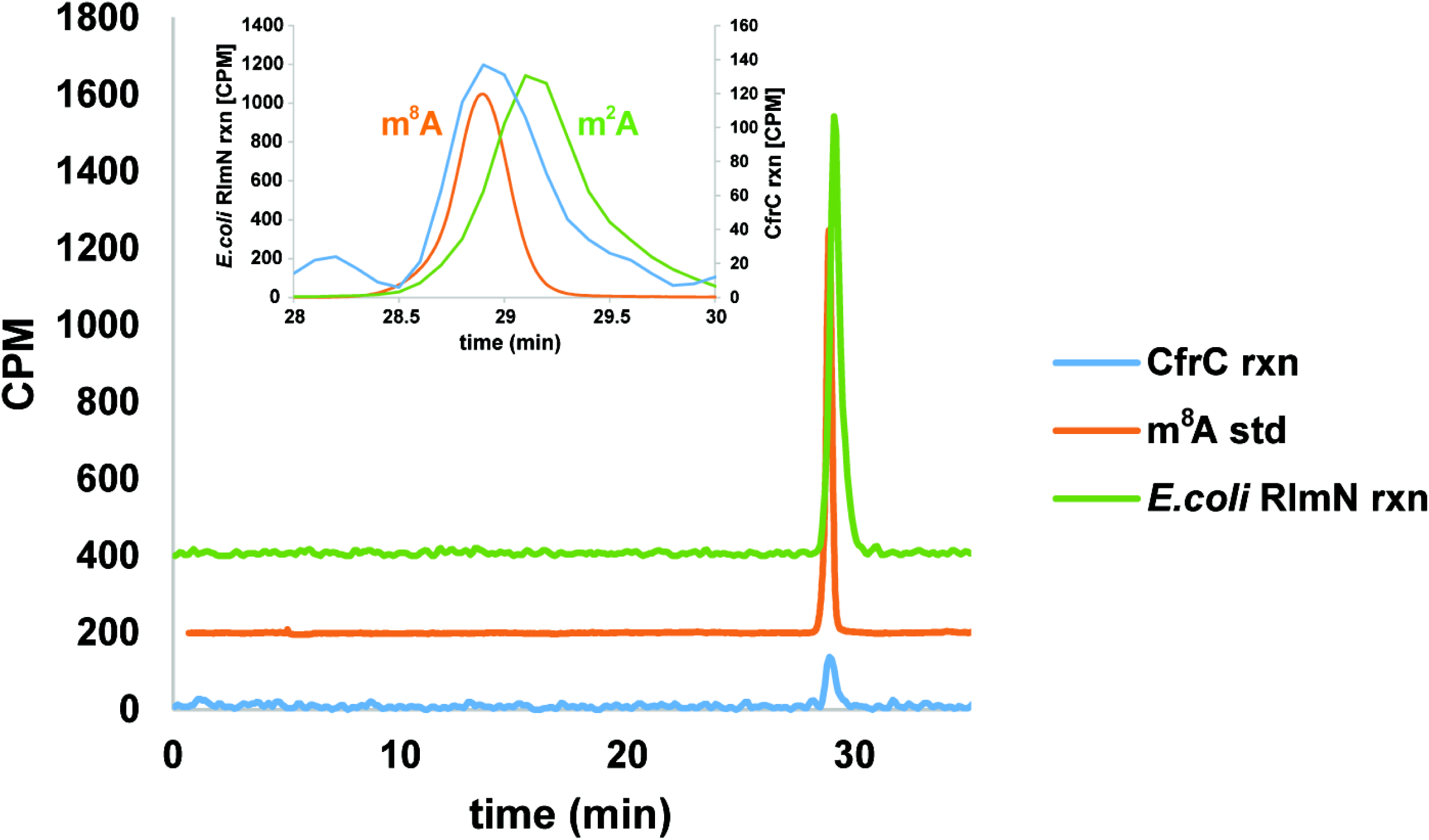
HPLC analysis of methylation products from Cfr(C) (blue) and *E. coli* RlmN reactions (m^2^A, green) with *E. coli* 2447-2625 rRNA fragment. A m^8^A standard is shown in orange.

## 4. Discussion

We investigated seven clinical *C. difficile* strains from Latin America that circulated between 2009 and 2016 to determine whether they carry functional *cfr* or *cfr*-like genes. Analysis of their draft WGS indicated the presence of various alleles of *cfr*-like genes, while phylogenetic analysis suggested the presence of a new Cfr-like clade comprised of Cfr(C) and a putative Cfr-like sequence.

We provide for the first-time *in vitro* evidence of the RNA methylation activity of Cfr(C). Combined with the observation of a PhLOPS_A_ phenotype in isolate DF11, the clustering of this novel putative *cfr*-like sequence with Cfr(C) suggests that its product could be a Cfr methylating enzyme implicated in antibiotic resistance. This hypothesis is yet to be experimentally verified.

The finding of *cfr*-like genes in various types of MGEs with partial hits to genomic sequences reported for other intestinal Firmicutes lends evidence for the plasticity of the *C. difficile* genome ^32^ and supports the role of this pathogen as a reservoir of resistance genes in the human gut ^33^. This situation is worrisome because linezolid is used for the treatment of methicillin-resistant *Staphylococcus aureus* ^34^ and vancomycin-resistant enterococci ^35^, which reside in the same Phylum as clostridial organisms. Indeed, versions of Tn*6218*, such as those detected in isolates HON06 and HON10, have been found among *Enterococcus faecium* isolates from German hospital patients ^36^.

The widespread detection of *cfr*-like genes among various epidemic NAP1/RT027/ST01 strains deserves attention to clarify whether this situation contributes to virulence. This notion is reinforced by the fact that linezolid and moxifloxacin resistance, a marker of highly virulent *C. difficile* strains, are often linked in this ribotype ^37^. Furthermore, since antibiotics are crucial both for the induction, progression, and treatment of CDI, multidrug-resistance (MDR) is particularly worrisome when present in epidemic types such as the NAP1/027/ST01 strain, which has been linked to severe disease and CDI outcomes ^38^.

Although the *cfr*(B) allele of isolates HON06, HON11, PUC51, and PUC347 is identical, the last two isolates did not show a strong PhLOPS_A_ phenotype. It has been shown that Cfr(B) is functional when encoded by Tn*6218* ^10,13^, hence we propose that this gene is not as active in PUC51 and PUC347 possibly due to genomic neighborhood effects or the lack of HTH transcriptional factors and RNA polymerase sigma factors seen in all other MGEs here reported. Proteins from these families commonly participate in gene expression regulation, and in *C. boltae* 90B3, *cfr*(C) is co-transcribed with the gene for a putative HTH DNA binding protein^12^.

To further support the role of *cfr*-like enzymes in antibiotic resistance, we have provided the first *in vitro* evidence that Cfr(C) methylates the C8 position in A2503 of *E. coli* 23S rRNA and in A2508 of *C. difficile* 23S rRNA. In this regard, the poor activity of Cfr(C) towards the assayed rRNA fragments could reflect differences in substrate requirements between Cfr(C) and *E. coli* RlmN ^26^ or result from the lack of other modifications in the RNA substrate that are necessary for efficient methylation by Cfr(C).

Our results likely reflect the unique patterns of antibiotic consumption that distinguish Latin America ^39^. Therefore, it would be worthwhile to analyze additional resistance phenotypes in strain collections from this region. This can be achieved through a combination of classical phenotypic tests, whole genome sequencing, and biochemical validation, as exemplified here. As already noted ^40^, a prompt phenotypic and genotypic identification of resistance genes, effective antimicrobial stewardship and infection control programs, and alternative therapies are needed to prevent and contain the spread of MDR *C. difficile* strains.

## Ethical Approval

None

